# RNA Atlas of Human Bacterial Pathogens Uncovers Stress Dynamics Linked to Infection

**DOI:** 10.1101/2020.06.29.177147

**Authors:** Kemal Avican, Jehad Aldahdooh, Matteo Togninalli, Jing Tang, Karsten M. Borgwardt, Mikael Rhen, Maria Fällman

## Abstract

Despite being genetically diverse, bacterial pathogens can adapt to similar stressful environments in human host, but how this diversity allows them to achive this is yet not fully understood. Knowledge gained through comparative genomics is insufficient as it lacks the level of gene expression reflecting gene usage. To fill this gap, we investigated the transcriptome of 32 diverse bacterial pathogens under 11 host related stress conditions. We revealed that diverse bacterial pathogens have common responses to similar stresses to a certain extent but mostly employ their unique repertoire with intersections between different stress responses. We also identified universal stress responders which shed light on the nature of antimicrobial targets. In addition, we found that known and unknown putative novel ncRNAs comprised a significant proportion of the responses. All the data is collected in PATHOgenex atlas, providing ample opportunities to discover novel players critical for virulence and maintenance of infection.

## Introduction

Bacterial pathogens with different genetic and physiological features share the capacity to sense and respond to external changes in the host via regulation of their global transcriptome. These responses, commonly called stress responses, are critical for pathogen invasiveness and protection from host immunity, which makes them topics of interest for future development of new antimicrobials. The responses are often complex, and synergistic regulation of regulatory networks can be pivotal in sensing and adapting to environments in different colonization niches during different phases of infection (Avican et al., 2015; Heine et al., 2018; Malachowa et al., 2015; Mandlik et al., 2011; Nuss et al., 2017). Infecting bacteria encounter numerous stress conditions which vary depending on the route and phase of infection. For many pathogens, one of the first changes upon infection of mammalian hosts is altered temperature. Consequently, bacterial virulence factors, such as type III secretion systems in *Yersinia* spp. and *Shigella* spp., are positively regulated by temperature (Bolin et al., 1988; Prosseda et al., 2004). Low pH in the gastrointestinal tract (GIT), genital tract, dental plaque, and skin, and acid exposure in phagosomes represents additional stressful environments that pathogenic bacteria have to adapt to (Lukacs et al., 1991; Lund et al., 2014). Another agent commonly affecting enteric bacteria are bile salts produced in the liver and secreted into the GIT, as well as secondary bile salts produced by the microbial flora (Hofmann and Hagey, 2008). The hyperosmotic nature of the blood stream and GIT can also be harsh for certain pathogens, inducing the expression of virulence genes in some cases, such as in *Helicobacter pylori* and *Vibrio cholerae* (Ishikawa et al., 2012; Loh et al., 2007). Limited nutrient, oxygen, and iron levels in the host are other stresses that most pathogens encounter and have to adapt to for survival (Fang et al., 2016). Furthermore, many invading bacteria encounter neutrophils and macrophages, which produce toxic reactive oxygen and nitrogen species.

Bacterial pathogens are diverse regarding gene content and regulatory networks, but yet equally successful to adapt to these host stresses. Comparative genomics studies have enabled accumulated knowledge of the diversity of bacterial pathogens. However, the question, how and when different gene products are employed by diverse pathogens to cope with same stresses encountered in human host remains to be answered. To complement comparative genomics studies and answer those questions, global gene expression profiling of diverse bacterial pathogens under host related conditions is desired. Even though there are large amount of gene expression data available for many bacterial pathogens, the collection of such dataset from different sources is not optimal to complement comparative genomics due to differences in sample handling, experimental methodologies, and incomplete datasets.

In this study, we analyzed the global expression profile of 32 different bacterial pathogens, representing 28 different species, under 11 infection-relevant stress conditions with same experimental set-up and methodology. We assessed expression profile of 105 088 genes and identified common and specific gene groups to certain bacterial groups with comparative genomics. All datasets were collected in an interactive RNA atlas, named PATHOgenex (www.pathogenex.org), freely available to the research community to interrogate expression profiles of human pathogens. The datasets were used to uncover similarities and discrepancies in different stress responses across different groups of bacteria. This was made possible by grouping genes present in different bacteria according to function and homology and computing a score showing the probability to be regulated in a particular environment. These scores also allowed us to identify universal stress responders, which are genes that are present in many bacterial species showing altered expression in multiple stress conditions. This group of genes could potentially serve as a source for studies aiming to identify novel targets of antimicrobials as it contain many already known targets. We could also show that non-coding regulatory RNAs can be differentially regulated in response to stressful environments and that novel small and long non-coding RNAs can be explored from the dataset.

## Results and Discussion

### Consensus In Vivo Stress Conditions

The majority of bacteria chosen for the PATHOgenex RNA atlas represent pathogens causing worldwide health problems. Most of the strains are commonly used in the microbial research community and diverse in terms of Gram staining, phylogeny, and oxygen requirement, which should facilitate interpretation and use of data **(Figure 1A)**. We designed 10 experimental setups mimicking different infection-relevant stress conditions expected to be encountered by bacterial pathogens in the human host: acidic stress, bile stress, low iron, hypoxia, nutritional downshift, nitrosative stress, osmotic stress, oxidative stress, high temperature and starvation (stationary phase). A very comprehensive suite of infection relevant conditions were previously described for *Salmonella enterica* serovar Typhimurium (Kroger et al., 2013). We used similar conditions that are applicable to diverse human bacterial pathogens. We also included exposure to specific *in vitro* virulence inducing conditions, such as shifting *Yersinia pseudotuberculosis* from 26°C to 37°C and depleting extracellular Ca^2+^, L-lactate supplementation for *Neisseria spp*. (Sigurlasdottir et al., 2017), and presence of charcoal-like resin XAD-4 in the growth medium for *Listeria monocytogenes* (Ermolaeva et al., 2004) **(Figure 1B)**. For some species, such as *Mycobacterium tuberculosis* and *Legionella pneumophila*, virulence inducing conditions are unknown and were not performed. As control for differential expression analyses, we utilized unexposed, exponentially grown bacteria. The growth temperature and culture medium were adjusted for each specific strain to achieve optimal growth conditions. Similarly, the strength of the stress with the different agents and associated exposure time was designed to be as similar and relevant as possible for each bacterial species. However, minor changes were necessary for certain conditions; for example, the “low pH level” used for *H. pylori* strains were lower than for the others, and a significantly higher H_2_O_2_ concentration was required to induce a stress response in *Acinetobacter baumannii* **(Table S1)**.

**Figure 1.**
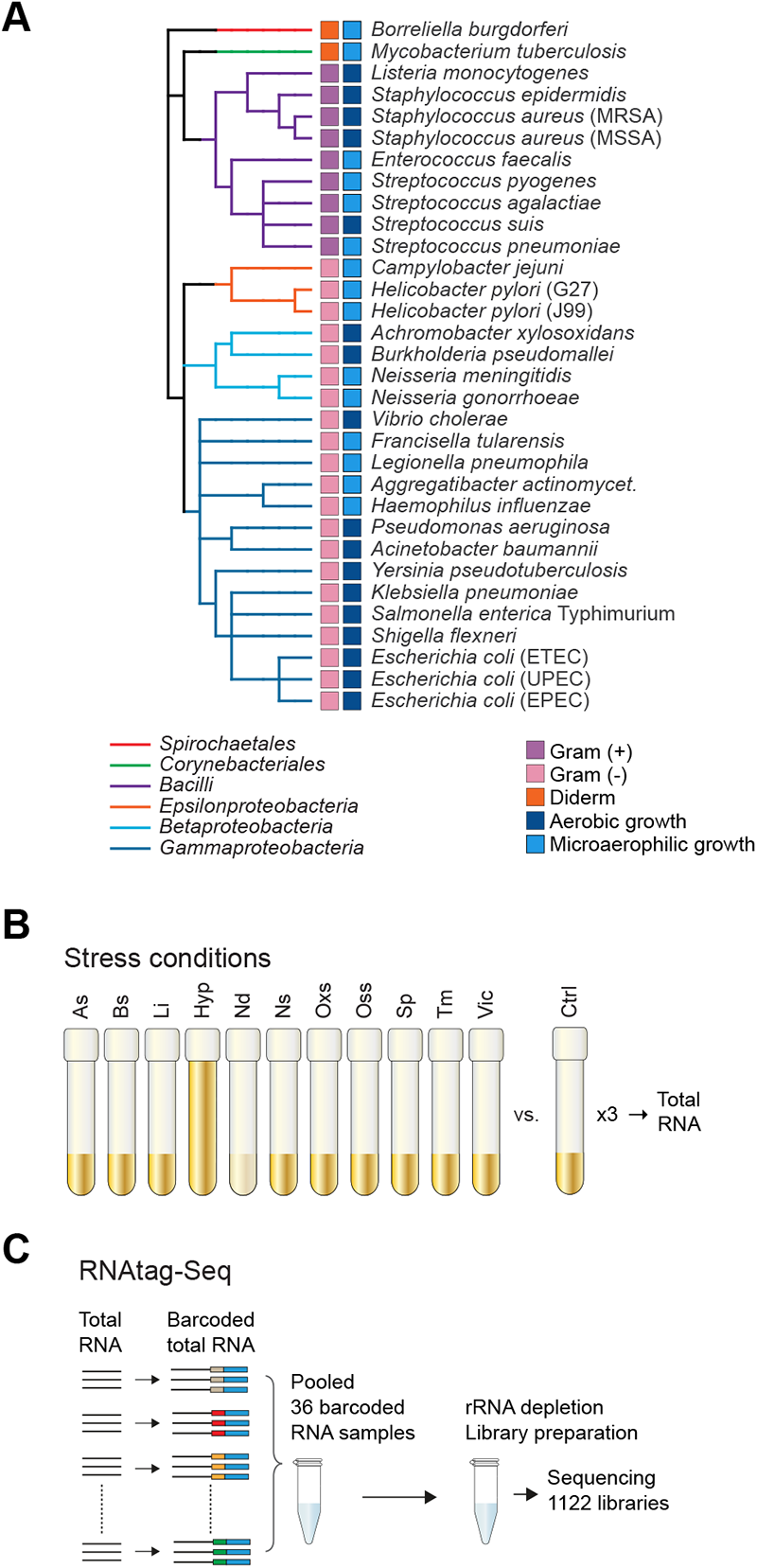
Experimental setup for PATHOgenex RNA atlas. **(A)** Phylogenetic clustering of bacterial species included in the study. Phylogenetic orders as well as Gram staining groups and oxygen dependency are indicated by color. **(B)** Schematic overview of the 11 infection-related stress conditions and control (Ctrl) sample not exposed to any stress: acidic stress (As), pH 3-5 for 10 minutes; bile stress (Bs), 0.5% bile salts for 10 minutes; low iron (Li), 250 μM 2,2-dipyridryl for 10 minutes; hypoxia (Hyp), low oxygen for 4 hours; nutritional downshift (Nd), incubation in 1X M9 salts for 30 minutes; nitrosative stress (Ns), 250 μM Spermine NONOate for 10 minutes; osmotic stress (Oss), 0.5% NaCl for 10 minutes; oxidative stress (Oxs), 0.5-10 mM H_2_O_2_ for 10 minutes; stationary phase (Sp); temperature (Tm), 41°C for 20 minutes; varied virulence inducing condition (Vic). Some adjustments were made for certain bacteria (see Table S1). **(C)** Schematic illustration of the RNAtag-Seq approach allowing multiple samples (36 in this study) per library used for obtaining 1 122 transcriptomes deposited in the PATHOgenex RNA atlas.

### Large Scale RNA-seq generates Accurate Datasets

All stress experiments were performed in laboratories with expertise in growing/studying the different strains **(Figure 1A** and **Figure S1)**. Three biological replicates were used for statistical analyses, in total 1 122 rRNA-depleted RNA samples were prepared using RNAtag-Seq, an approach based on tagging RNA samples with barcode oligonucleotides, enabling pooling of multiple samples in one RNA-seq library (Shishkin et al., 2015) **(Figure 1C).** This strategy is expected to limit biases during sample preparation. We used a set of 36 previously described barcode DNA oligonucleotides (Shishkin et al., 2015) in which the same set of three oligos was applied for each condition for consistency **(Key Resources).** The sequencing reads from each library were demultiplexed according to the 8-nt unique barcode reads to separate the different samples. The number of reads for the different barcoded samples did not reveal significant variation across conditions and species **(Table S2),** excluding bias regarding over-representation of one or a set of barcodes during library preparation. We obtained an average of 12.2 million reads for each rRNA-depleted barcoded library, which far exceeds the 2-3 million reads considered sufficient to determine differentially expressed genes from bacteria with high significance (Haas et al., 2012). The average mapping of the reads to the reference genomes was >90% **(Table S2, Key Resources)**. The lowest coverage (67.3%) was seen for the clinical isolate of *Achromobacter xylosoxidans*, likely due to the lack of an appropriate reference genome. In this case, we used the sequenced *A. xylosoxidans* genome that gave the highest number of mapped reads. In general, the read mapping to reference genomes showed efficient rRNA depletion, with only 0.1-5% reads mapping to rRNA and tRNAs, except for *Campylobacter jejuni*, two of the *H. pylori* strains, and *Neisseria gonorrhoeae*, which had higher proportions mapping to rRNA **(Table S2)**. Nevertheless, the library sequencing for these species was deep enough to cover 94% of the coding sequences (CDSs), with >10 reads per CDS in the least efficiently depleted library.

### Genome Size and Infection Niche Influence Gene Expression

To determine what proportions of genes are active or silent under tested conditions we calculated the expression levels of annotated CDSs, the transcript per million (TPM) values, for all strains. Hierarchical clustering of expression profiles clustered replicates of the same sample together, indicating a robust measurement of gene expression in all conditions and strains **(Figure S2)**. For gene expression analysis, genes with TPM ≥ 10 were considered to be active and genes with TPM < 10 were considered to be silent, which is similar to definitions used by others (Kroger et al., 2013; Wagner et al., 2013). We found that 32-95% of all genes in each of the 32 strains were constitutively active across the 12 tested conditions. There were also genes only active in certain conditions (one to 11 conditions), but a set of genes (0.7-20%) were silent in all of the conditions **(Figure 2A).** The silent genes may require more complex environments with synergistic effects of many stresses, as in real *in vivo* environments, to be expressed. It was obvious that bacterial species with relatively smaller genomes had a greater proportion of active genes in all conditions **(Figure 2B)**. This may indicate conservation of indispensable genes during the evolutionary shrinkage of the genomes. Reductive evolution is a well-known event in endosymbionts and pathogenic bacteria with respect to adaptation in defined niches (Miravet-Verde et al., 2017). This is illustrated by the larger genomes of ubiquitous environmental species, such as *E. coli*, and smaller genomes of niched pathogens, such as *Borrelia burgdorferi*. A more flexible response was seen in turning transcription on and off in bacteria with larger genomes, which may be related to the function and number of regulatory proteins within the species. Comparative genomics studies of gene content and transcription machinery have previously demonstrated that the number of transcription factors per open reading frame increases with genome size in many pathogenic bacteria (Perez-Rueda et al., 2009). We observed the same pattern within the set of pathogens in this study **(Figure 2C)**. Thus, previous assumptions of less flexibility in turning gene expression on and off in pathogens with reduced genomes were here verified at the RNA level across multiple bacterial species.

**Figure 2.**
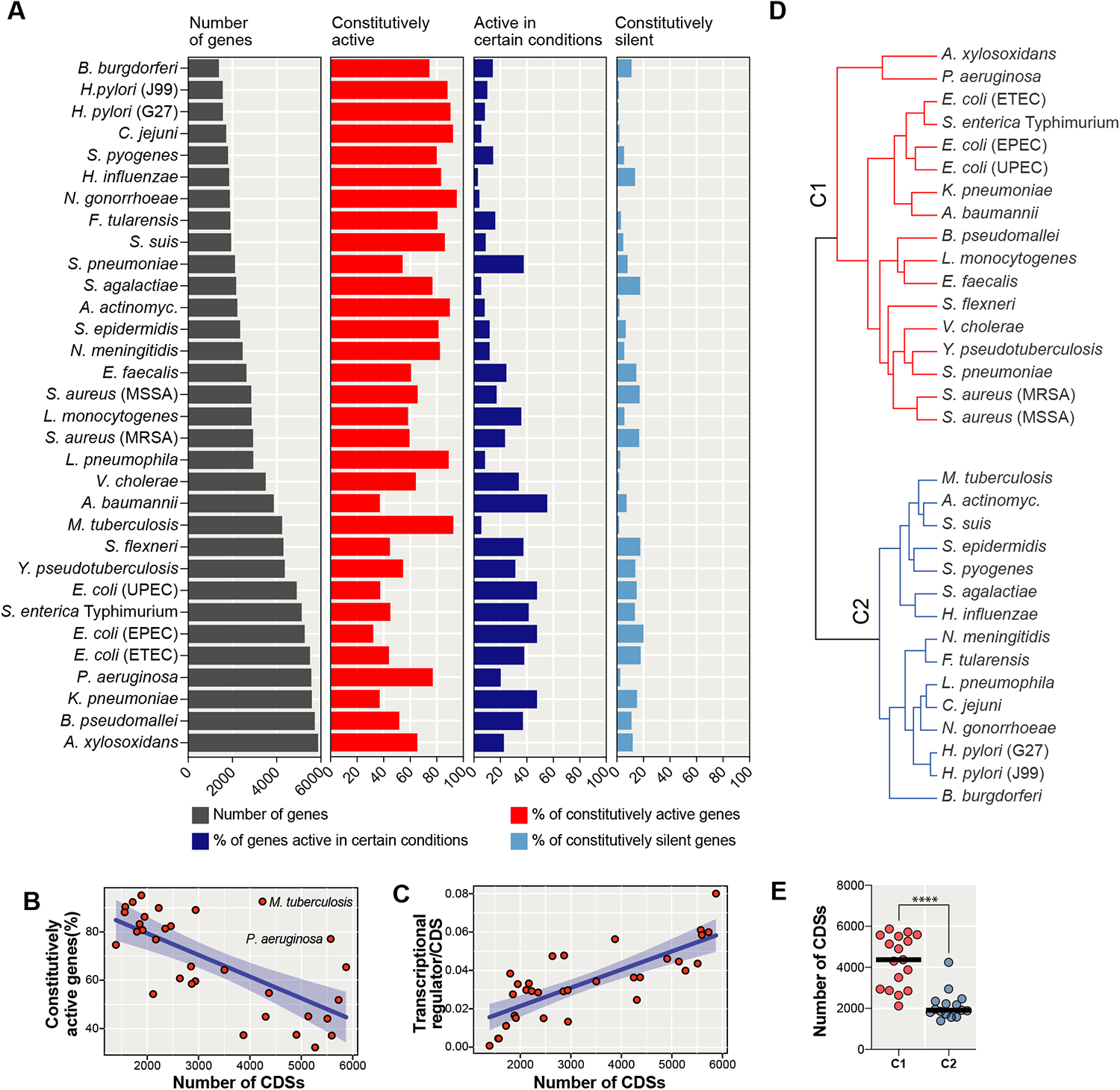
Niched pathogens have high proportion of constitutively active genes. **(A)** Diagrams showing; number of genes (gray), percentage of constitutively active genes (red), percentage of genes active in certain (one to 11) conditions, and percentage of constitutively silent genes. Genes with TPM≥10 are designated as active, and TPM<10 as silent. Bacterial species are sorted according to the number of CDSs in their reference genome. **(B)** Scatter plot showing percentage of constitutively active genes for each species. **(C)** Scatter plot showing the number genes annotated as transcriptional regulator per CDS for each species. Blue trend lines in (B) and (C) show the degree of linear regression with a 95% confidence interval. **(D)** Hierarchical clustering of all strains based on global gene expression profiles in (A). The 2 main clusters C1 (red) and C2 (blue) are indicated. **(E)** Dot plot showing the number of CDSs in C1 (red) and C2 (blue),. The significance between the two groups was calculated using unpaired t-test. ****P < 0.0001.

Most of the bacteria with reduced genomes represent pathogens with restricted niches, which match the low requirement of dynamic changes in gene expression **(Figure 2A)**. *M. tuberculosis* and *P. aeruginosa* did not follow this trend (**Figure 2B)**, suggesting that reduced regulation of gene expression is not always due to genome reduction. Hierarchical clustering based on profiles of constitutively active genes, genes active in certain conditions, and silent genes generated two bacterial clusters: ubiquitous environmental species (C1) and niched pathogens (C2) **(Figure 2D).** Comparing the genome sizes of the strains in the two clusters clearly showed that the strains with restricted infection niches had smaller genomes **(Figure 2E)**. Therefore, our data verify at the level of RNA expression that bacterial pathogens with restricted niches, mostly with reduced genomes, have lower plasticity to turn gene expression off.

### Bacteria Respond to Stress via Dynamic Changes in Gene Expression

To reveal gene expression patterns associated with responses to different environmental stimuli we performed differential gene expression analyses for each strain. Compared to gene expression in unexposed, exponentially grown control bacteria, most bacteria dynamically responded to the stresses by regulating 62-90% of their genes in at least one condition **(Figure 3A).** The highest fraction of regulated genes was measured under hypoxia, nutritional downshift, and stationary phase conditions **(Figure 3B)**. The differential expression analysis was verified by checking for differential expression of known genes that are expected to be regulated under certain stress conditions. We found that most genes expected to be induced under the different conditions were upregulated in the species harboring the gene, clearly showing the accuracy of the data **(Figure S3A).**

**Figure 3.**
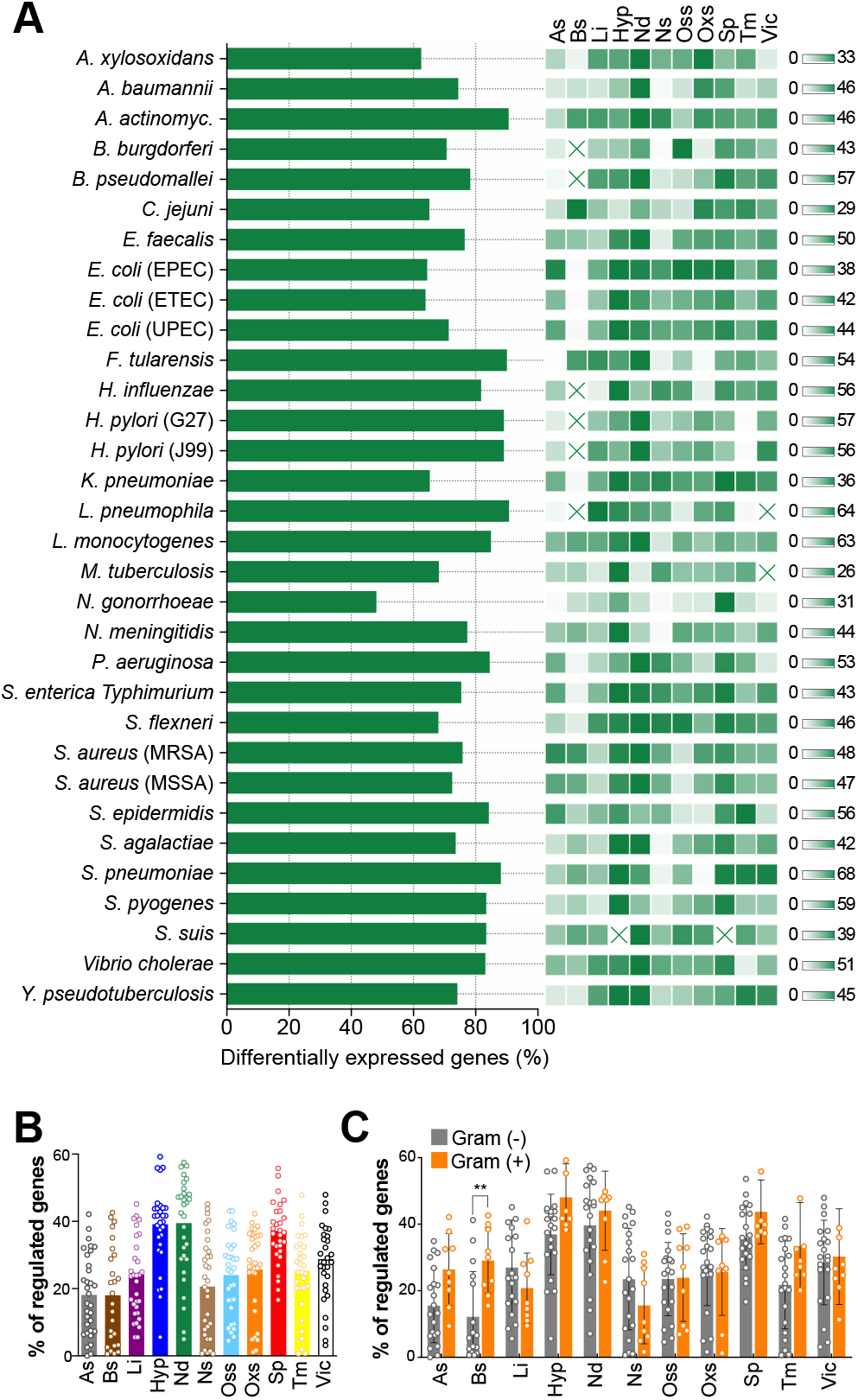
Cross-Microbial Human Pathogens Dynamically Respond to Changes in the Environment. **(A)** Proportion of genes differentially regulated in at least one condition for each bacteria. The linked heat maps show proportions of regulated genes for each stress condition. The scales to the right show zero (white) to highest (green) proportion for each species. Heat map squares marked with a cross indicate that RNA-seq was not performed for that specific condition. **(B)** Dot plot showing the percentages of genes differentially regulated in each condition for all included species. The bar plot show the mean value for each condition. **(C)** Dot plot showing percentages of differentially regulated genes in each condition for Gram-negative (gray) and -positive bacteria (orange). The bar plots show the mean value for each condition. The significance between the groups was calculated with Multiple t-test using Holm-Sidak method. * indicates adjusted *p*-value < 0.05.

Given that different bacteria have distinct features, we also analyzed the levels of responses for different groups of bacteria, such as Gram-negative and -positive, aerobic and microaerophilic, different phylogenic orders **(Figure 1A),** and the niched pathogens versus environmental species groups **(Figure 2D).** The levels of responses were relatively similar between different groups. For Gram-negative and -positive strains, however, responses to bile were significantly higher in the latter group, possibly reflecting a higher magnitude of stress due to having only one plasma membrane **(Figure 3C, Figure S3B, C, and D).**

### Clustering Genes into Gene Groups for Cross Microbial Comparisons

Comparison of differential expression of genes with similar functions or homologous genes across different bacterial species is important for a broad and comprehensive understanding of gene regulation and gene function in terms of adaptation to new environments. Therefore, we clustered genes from the 32 strains into gene groups using two different orthology and homology approaches: one based on functional orthologs using KEGG’s annotation tool GhostKOALA (Kanehisa et al., 2016), and one based on isofunctional homologs using the PATRIC database with Patric Global Family (PGFam) (Wattam et al., 2014). KEGG orthology (KO) numbers are assigned to genes harboring the same function, independent of sequence homology, and can include genes with different sequences and lengths. For example, KO0123 is assigned to the genes encoding formate dehydrogenase major subunit, which includes all major subunits of formate dehydrogenase, including FdhF, FdnG, FdoG, and NapA in our dataset. On the other hand, PGFam numbers are assigned to genes with the same function and sequence homology (i.e., isofunctional homologs) as indicated by RAST and CoreSEED (Davis et al., 2016). Therefore, PGFam groups are more specific in terms of function and homology, but with fewer genes per group. Consequently, in contrast to KO groups, formate dehydrogenase subunits were assigned to four different PGFam groups. For KEGG, the tool assigned 6054 KO numbers to 63 831 genes, representing 60.7% of the total 105 088 genes in the dataset **(Figure 4A)**. For PGFam, 27 999 PGFam numbers were assigned to 97 565 genes, representing 92.8% of the total number of genes in the dataset.

**Figure 4.**
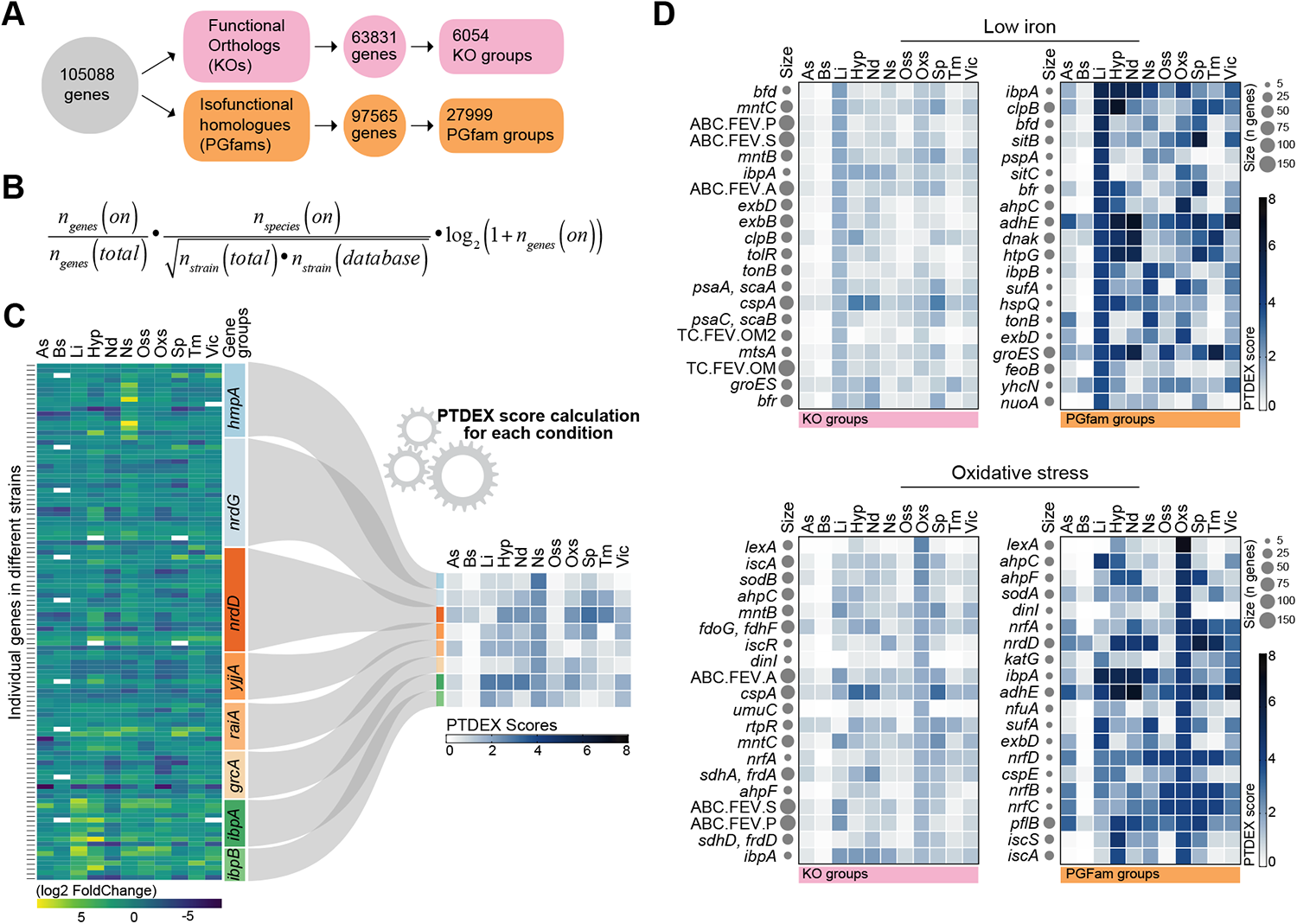
Transformation of differential expression to PTDEX scores enables comprehensive understanding of gene regulation under stress conditions. **(A)** Schematic illustration of clustering of 105 088 genes based on functional orthology (KO) and isofunctional homology (PGFam) and resulting KO and PGFam gene groups. **(B)** Equation used to calculate PTDEX scores for KO and PGFam gene groups. **(C)** Illustration of the transformation of differential expression values of genes from many pathogens that are clustered in one gene group into a PTDEX score for each stress condition. Eight gene groups with highest PTDEX scores in nitrosative stress condition are shown as example. White rectangles indicate no differential expression. **(D)** The 20 KO and PGFam groups with highest PTDEX scores in low iron (top) and oxidative stress (bottom) conditions. The size of gray dots after the gene name relates to the number of genes in the gene groups.

### Revealing Probability of Gene Groups to be Differentially Expressed

To predict the probability of the genes within the KO or PGFam groups to be regulated under certain stress conditions, we formulated an equation that computes a stress condition-specific score indicating differential regulation of genes in the group, defined as ‘<u>p</u>robability <u>t</u>o be <u>d</u>ifferentially <u>e</u>xpressed (PTDEX) score’. The equation was employed to gene groups that contain at least 2 genes (5 340 KO groups and 11 353 PGFam groups) **(Table S3 and S4)**. The resulting PTDEX score takes into account how well conserved the genes in the group are among the 32 pathogens, what proportion of the genes in the group are differentially regulated under a particular stress condition, and how well this regulation is preserved among the 32 strains **(Figure 4B)**. For example, a gene group that is present in all 32 species and differentially expressed under acidic stress in many of the species will have a high PTDEX score for acidic conditions. This not only indicates a high probability of the genes in this group being differentially expressed under acidic stress, but can also hint regulation of homologous genes from bacterial pathogens not included in this data set. Hence, using PTDEX scores for comprehensive understanding of gene regulation in bacteria is a novel approach providing in depth overview of common and specific regulation of same genes in different species and simplifies cross microbial comparisons **(Figure 4C)**.

To test the power of the resulting PTDEX scores, we ranked the KO and PGFam groups based on PTDEX scores in low iron and oxidative stress conditions and plotted the groups representing the highest 20 **(Figure 4D and 4E)**. Both KO and PGFam PTDEX scores were found to be indicative for regulation under these conditions. Many of the genes with high PTDEX scores for low iron represented genes encoding proteins known to be involved in iron uptake and iron homeostasis, such as *tonB, exbB, exbD, mntABC, bfr, bfd, ibpA, ibpB, sufA, feoB*, and *nuoA* **(Figure 4D)**. Similarly, the genes with highest KO and PGFam PTDEX scores in oxidative stress were genes known to respond to DNA damage and oxidative stress, such as *lexA, soda, sodB, aphC, dinI, umuC, rtpR*, and *iscA* **(Figure 4E)**. As expected the highest 20 were not exactly the same for the KO and PGFam groupings, but most of the expected gene groups were found in both. The higher PTDEX score observed for PGFam groups is likely due to more similar regulation of isofunctional homologues than of functional orthologs. Therefore, to derive more accurate conclusions, we used PGFam PTDEX scores for the remainder of our analysis.

### PTDEX Scores Enable Comparisons Between Stress Responses of Evolutionary Distinct Bacterial Groups

The gene expression data in PATHOgenex were collected from diverse bacterial species **(Figure 1A)** with distinctive gene content and transcriptional regulatory networks. Common transcriptional regulatory patterns, between for example Gramnegative and -positive bacteria, are anticipated to be rare. Therefore, we also reclustered genes of Gram-negative (21 strains from *Gammaproteobacteria, Betaproteobacteria*, and *Epsilonproteobacteria*) and Gram-positive strains (9 strains from *Bacilli*) into G- and G+ PGFam groups and then re-computed the PTDEX scores separately **(Table S5** and **S6).** We then generated a similarity matrix with the Pearson correlation coefficient for PTDEX scores from 7105 G-PGFam groups representing 47 468 genes and 2903 G+ PGFam groups representing 14 200 genes for each stress condition. It was here obvious that despite the substantial difference in PGFam groups **(Figure 5A)**, parts of the responses to external changes were indeed shared by Gramnegative and -positive bacteria **(Figure 5B)**. The conditions that showed highest overlaps were hypoxia, nutritional, and stationary phase.

**Figure 5.**
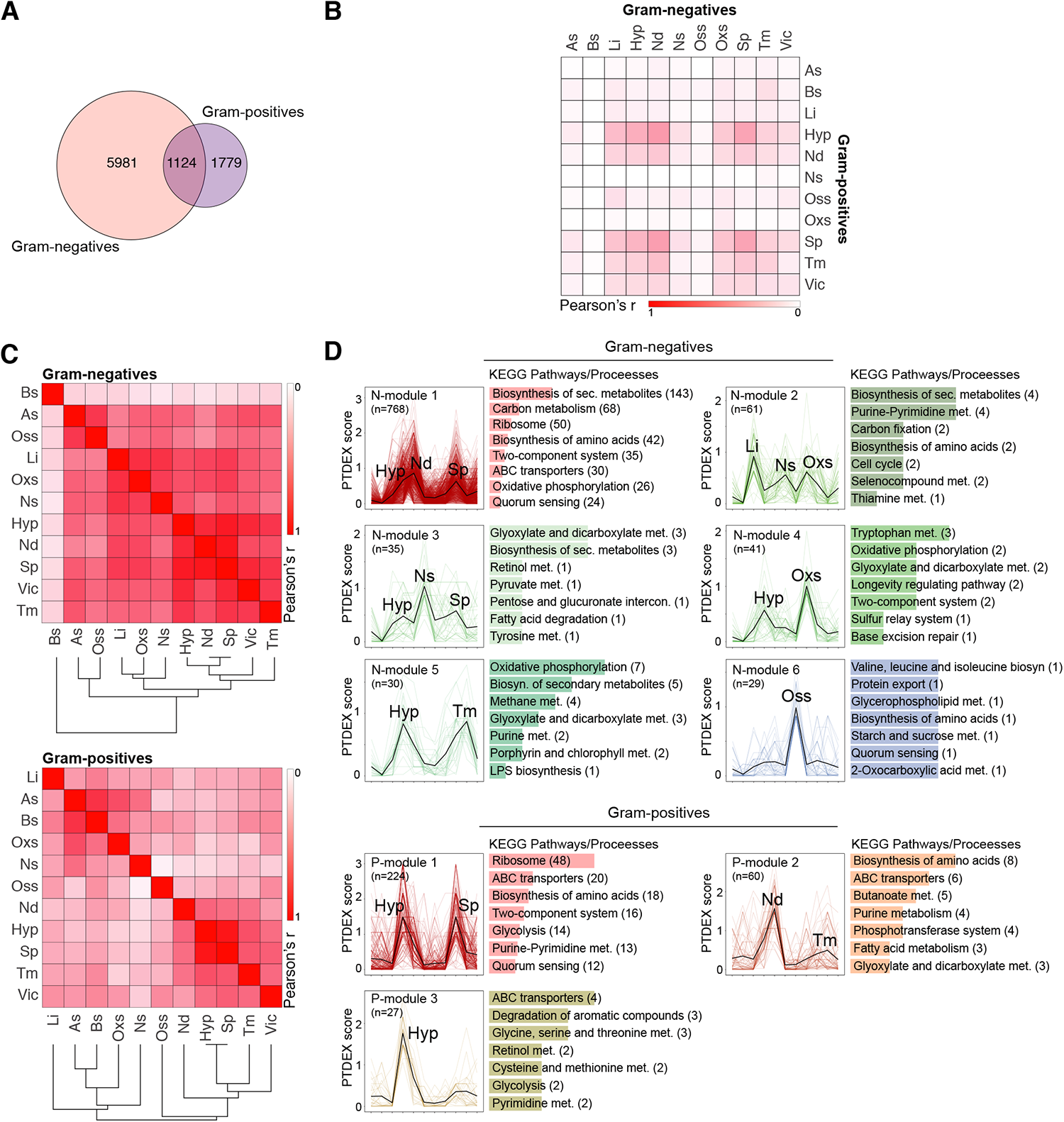
Gram-negative and -positive bacteria exhibit distinct responses. **(A)** Venn diagram of individual and shared PGFam groups with a PTDEX score >0 in at least one of the stress conditions. **(B)** Heat map showing degree of similarity between responses to different stresses in Gram-negative and -positive bacteria. Similarity was calculated using the Pearson correlation coefficient of PGFam groups’ PTDEX scores in each condition. **(C)** Heat map showing degree of similarity between responses to different stress conditions in Gram-negative and -positive bacteria. Similarities were calculated as in (B) and the similarity distances shown in a dendrogram. **(D)** Modules generated by CemiTool showing conditions that involve similar regulation of gene groups with high PTDEX scores in Gram-negative and -positive bacteria. *n* indicates number of PGFam groups within each module. The most enriched KEGG pathways/processes for each module are shown with the number of gene groups indicated in brackets. See also Table S5 and S6.

Despite this similarities there is significant differences in responses by Gram-positive versus Gram-negative strains, which supports previous studies (Rodionov, 2007). Transcription factors, especially global regulators, are poorly conserved in general, and transcriptional regulation has been found to be more flexible than their target genes (Lozada-Chavez et al., 2006). In support of this, an analysis of differential expression of transcription factors in response to the different stresses revealed high diversity between Gram-negative and Gram-positive strains, as expected (**Figure S4**). Most of the transcription factors were unique to either Gram-positive or -negative bacteria, and the few homologs present were mostly regulated differently. Further, the observed difference in response to bile, where Gram-positive, but not Gram-negative bacteria responded with expensive differential gene expression (**Figure 3C**) was also reflected here, with very few high PTDEX score transcription factors for these conditions in Gram negatives **(Figure S4).**

### PTDEX Scores Enable Identification of Intersections Between Stress Responses

Reponses to different stressors are expected to partly overlap due to involvement of the same molecular pathways, as well as functional diversity of gene products. Many stress responses are for example expected to involve halted growth, enabling translational adjustments and redirection of resources. To identify overlaps between different stress responses, the PTDEX scores of G-/G+ PGFam groups were used to compare responses among the Gram-negative and -positive bacteria separately. These comparisons indicated a higher degree of overlap between responses in Gramnegative bacteria, where especially hypoxia, stationary phase, and nutritional downshift overlapped to a high extent. Furthermore, also responses to low iron, oxidative and nitrosative stress, as well as responses to acidic and osmotic stress overlapped **(Figure 5C)**.

To reveal overlapping pathways and processes, we next implemented a co-expression module identification algorithm (Russo et al., 2018) on G- and G+ PGFam PTDEX scores. This algorithm generated six modules of PGFam groups for Gram-negatives showing common patterns of PTDEX scores in certain combinations of stress conditions **(Figure 5D, Table S5).** KEGG pathway mapping of the genes in these modules indicated overlapping pathways and processes involved in different stress responses **(Figure 5D**, N-module 1-6). It was obvious that the response to hypoxia partly overlapped with responses from all of the other stress conditions (N-module 1 and 3-5), except the response to osmotic stress that seemed to be unique (N-module 6). The largest overlap was seen between responses to hypoxia, nutritional downshift and stationary phase, reflecting the complexity and high requirement of multiple pathways for managing these situations. The overlapping expression patterns included many genes involved in carbon metabolism, protein synthesis, oxidative phosphorylation and quorum sensing (N-module 1). Responses to low iron, nitrosative, and oxidative stress also shared PTDEX score patterns (N-module 2). In Gram-positive strains, the degree of overlap between stress responses was lower **(Figure 5C)** and only three modules were generated **(Figure 5D**, P-module 1-3). The largest overlap here was between hypoxia and stationary phase (P-module 1), and to a lesser extent between nutritional downshift and temperature (P-module 2). In contrast to that seen for Gram-negative strains, some PGFams groups with high PTDEX scores were specific to hypoxia only (**Figure 5D**, P-module 3).

### PTDEX Scores Reveal Universal Stress Responders Including Known Antimicrobial Targets

Although the Gram-negative and -positive bacterial pathogens studied here differ in gene content, the preserved genes may have fundamental roles for bacterial physiology and adaptation to stressful environments. Participation in multiple stresses can be key to their conservation during the evolutionary diversification of species. To retrieve genes encoding these universal stress responders (USRs), we selected the PGFam groups that were differentially expressed in at least six conditions and contained at least 11 genes from Gram-negative strains (21 strains in total) and 4 genes from Gram-positive strains (9 strains in total) **(Table S7)**. We used the general PGFam PTDEX scores and the threshold for *‘high PTDEX score’* was determined to be ≥0.25, which indicated the score value where at least 50% of the genes in the group are differentially expressed **(Figure S5)**. These criteria allowed identification of 421 USRs. Functional clustering of the USRs with PATRIC subsystems (Wattam et al., 2017) revealed that USRs are involved basic biological processes **(Figure S6)**. Interestingly, 24 of the identified USRs were genes representing targets for antibiotics, where mutations in their sequences been shown to confer antibiotic resistance **(Table 1)**. Hence, it is thus likely that novel putative antibiotic targets are to be found among remaining 397 USRs.

**Table 1.**
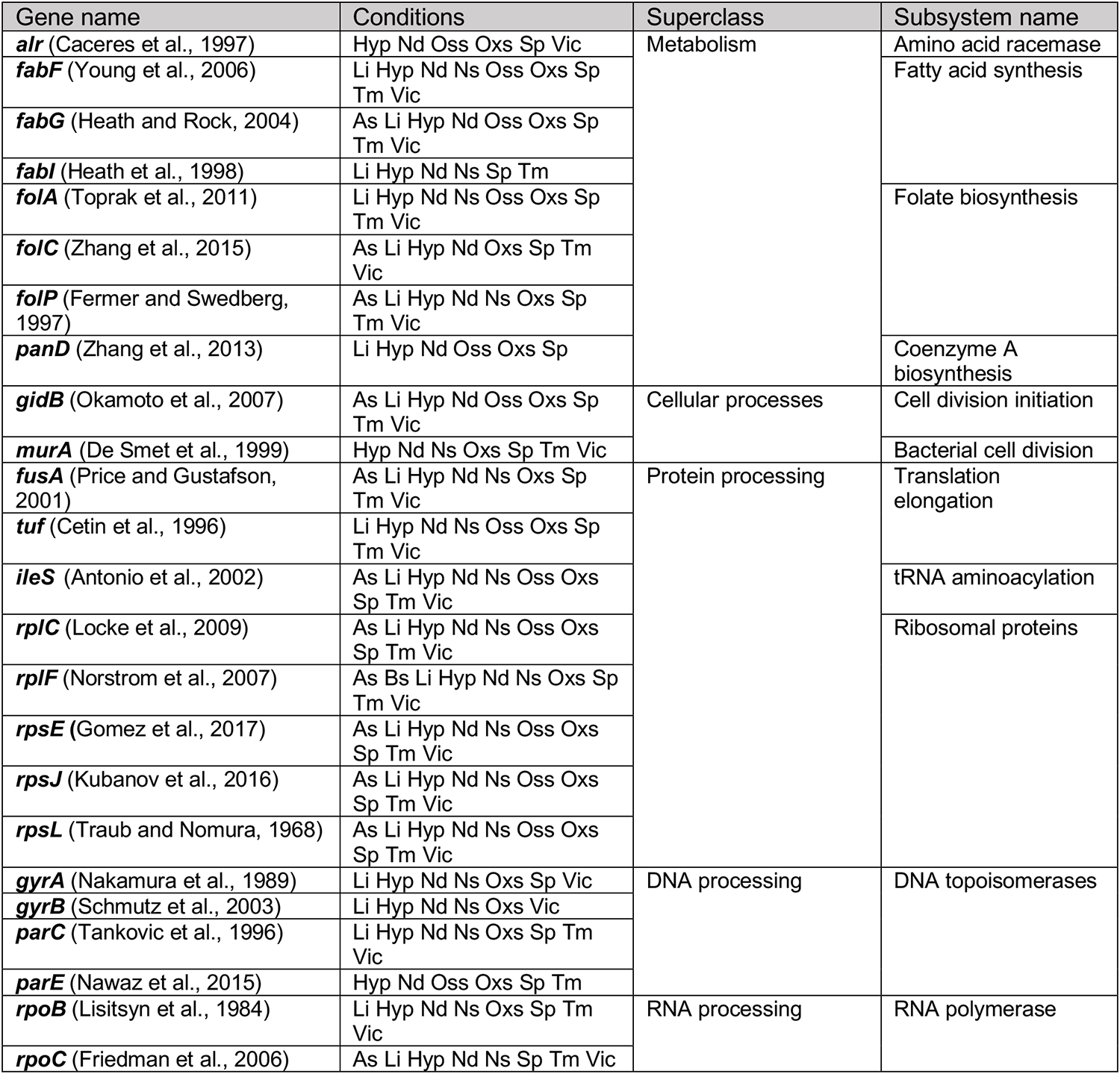
USRs known to be target for antimicrobials

### Exploring Stress Related Expression from Non-coding Regions

In addition to differential expression of annotated genes, this data set also provides information of the complete transcriptional landscapes in different pathogens. Analysing overall transcription, we surprisingly observed that a substantial proportion of RNAs were transcribed from non-coding DNA regions, including sRNAs, regulatory non-coding RNAs, 5’ untranslated regions (UTRs), 3’-UTRs, and intergenic regions **(Table S2)**. The proportion is expected to be even higher than calculated here since the majority of sRNAs were excluded during sample preparation due to the >100 nt cut-off for library preparation. In some bacteria, expression of non-CDSs was higher than CDSs in hypoxia, nutritional downshift, and stationary phase conditions. Such a large number of transcripts from non-CDSs in bacteria has not been reported previously, likely due to studies focusing on CDSs usually only consider annotated CDSs for mapping reads, and that studies of non-coding RNAs generally use RNA purification methods dedicated to sRNAs. Noteworthy, for most species, the percent of transcripts from non-CDS was increased upon stress exposure in comparison to control **(Figure 6A).** This indicates that expression from non-coding regions also contribute bacterial response to stress conditions.

**Figure 6.**
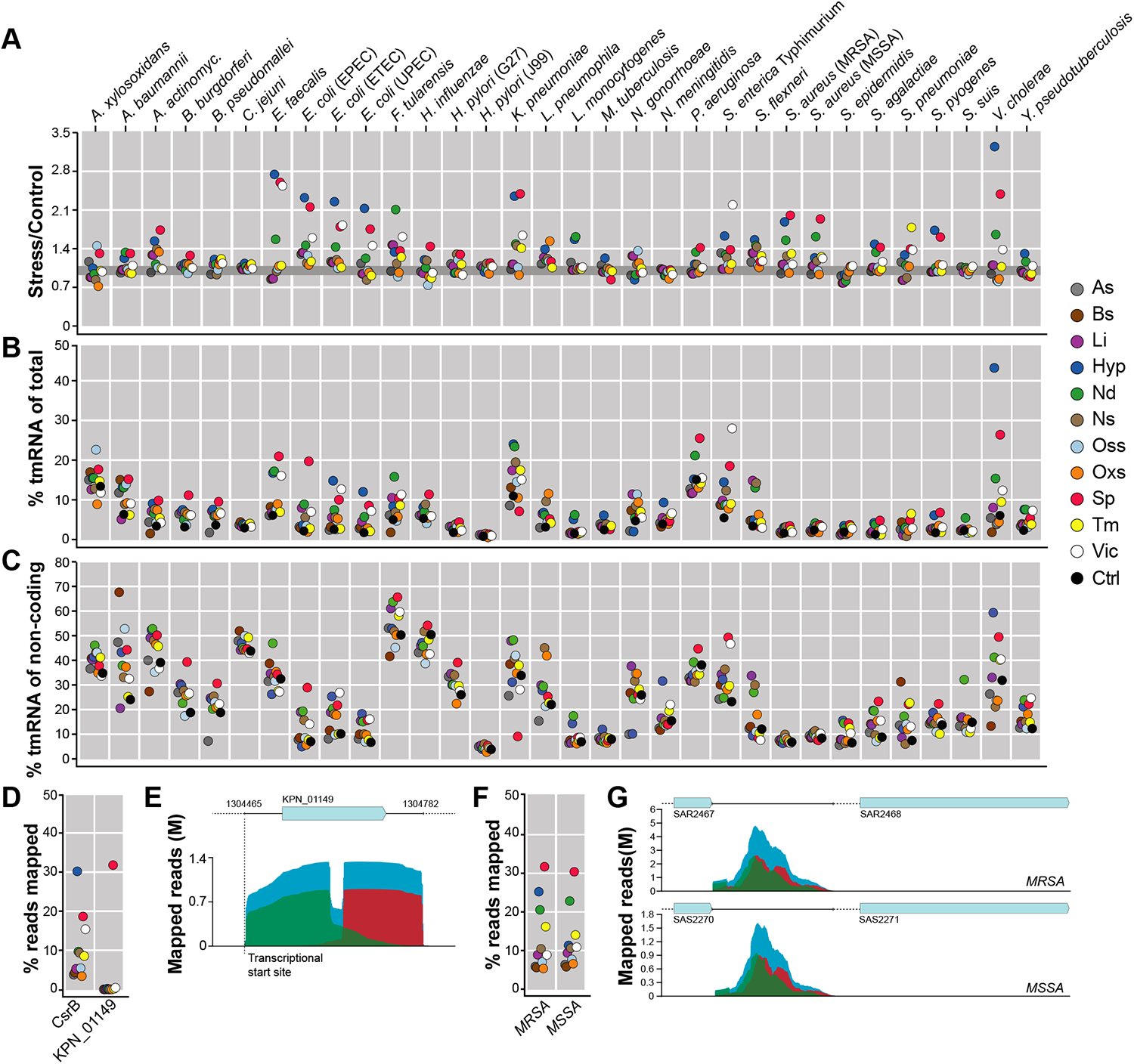
Complex stress conditions commonly involve differential expression from non coding regions, including tmRNA and novel non-coding RNAs. **(A)** Ratio of percent of reads mapped to non-CDSs under each stress condition in comparison to control for all tested strains. Different stress conditions are indicated by color. The gray line indicates Stress/Control value equal to one. **(B)** Proportion of reads mapped to tmRNA sequences of total number of mapped reads under each stress condition for all tested strains. **(C)** Proportion of reads mapped to tmRNA sequences of total number of reads mapped to non-CDSs under each stress condition for all tested strains. **(D)** Proportion of reads mapped to the *K. pneumoniae* CsrB and to the region covering KPN_01149 CDS with 5’-UTR and 3’-UTR sequences under each stress condition. **(E)** Number and aligments of reads mapped to KPN_01149 and its UTRs in stationary phase. Green indicates forward reads, red indicates reverse reads, and blue indicates paired reads. The vertical dashed line indicates the position of the TSS of KPN_01149 (Kim et al., 2012). **(F)** Proportion of reads mapped to the 1230-nt intergenic region in *S. aureus* MRSA and MSSA genomes under each stress condition. **(G)** Number of reads mapping to the 1230-nt region in *S. aureus* MRSA and MSSA strains in stationary phase. Colors represent reads as noted in (E). Different stress conditions are indicated with colored dots on the right. The means of three replicates in each condition were used for the calculation in (A), (B), (C), (D), and (F).

### Stress Responses Includes tmRNA and Various non-coding RNAs

To reveal the non-CDS transcripts responsible for the observed increases of non-CDS transcripts we analyzed the read mappings and found that the stress associated increases in many cases correlated with increased thexpression of tmRNA **(Figure 6B).** This is interesting as tmRNA, also known as SsrA, is a conserved and abundant type of RNA in bacteria that is important for trans-translation, a process that restores translation in detrimental situations (Gueneau de Novoa and Williams, 2004; Keiler, 2007; Moore and Sauer, 2005). For many bacteria, lack of tmRNA results in growth defects, sensitivity to various stress conditions, and attenuation of virulence (Brito et al., 2016; Munavar et al., 2005; Okan et al., 2006; Svetlanov et al., 2012). The increased expression of tmRNA **(Figure 6B, Table S8)** indicates that trans-translation is part of a common adaptation to challenging environments. The induction of tmRNA are particularly high in stationary phase, hypoxia, and upon nutritional downshift, which indeed are conditions associated with halted growth. In line with this, *smpB*, encoding a protein that interacts with tmRNA, was here identified as a USR with high PTDEX scores in stationary phase, hypoxia, and nutritional downshift **(Figure S6)**.

The level of tmRNA was however not high for all strains and conditions **(Figure 6C),** suggesting that also other non-coding RNAs are induced upon external stress. This was for example the case for *Klebsiella pneumoniae* (**Figure 6B)**, where an analysis of the mapped reads across its genome revealed that expression of the carbon storage regulatory non-coding RNA CsrB correlated with expression ratios for the intergenic regions in many the stress conditions **(Figure 6D)**. CsrB in *Klebsiella* spp. has not been extensively investigated, but has in other species been shown to be involved in regulation of many biological processes, such as carbon metabolism, biofilm formation, quorum sensing, and virulence (Fortune et al., 2006; Lenz et al., 2005; Liu et al., 1997). However, even though expression of CsrB correlated with increased transcription of non-CDSs in most of the conditions, this was not the case for stationary phase (**Figure 6D).** For the stationary phase we found high extent read mapping to 5’-UTR and 3’-UTR regions of a CDS (KPN_01149) encoding a hypothetical protein **(Table S2, Figure 6D and E).** The expressed region started at a position previously reported to be a transcriptional start site (Kim et al., 2012), supporting the accuracy of this finding (**Figure 6E**). The function of KPN_01149 is not known, but the presence of stress-induced bacterial acidophilic repeat motifs supports a role in stress responses. Similar to that of *K. pneumoniae* also *S. aureus* showed expression from non-coding regions that not corresponded with expression of tmRNA. Here we found high level of reads mapping to a 1230-nt intergenic region situated on the negative strand between two CDSs encoding a hypothetical protein and an AraC family transcriptional regulator **(Figure 6F and G**). Interestingly, by analysing data from a previous study (Szafranska et al., 2014) we found that expression of this intergenic region is increased during *S. aureus* infection **(Figure S7B)**. This sequence is only found in *S. aureus* and two other *Staphylococcus* spp., *S. argenteus* and *S. schweitzeri*. Though *S. argenteus* is known to colonize humans, *S. schweitzeri* isolates have been recovered from fruit bats and non-human primates, including gorilla (Becker et al., 2019). Given the broad repertoire of different environmental conditions and the number of species, detailed analyses of mapped reads in the different data sets provide opportunities for finding novel intergenic non-coding RNAs or UTRs. Both *K. pneumoniae* and *S. aureus* are examples of human pathogens responsible for causing troublesome infections, often involving biofilm and long-term treatments, thereby contributing to increased antibiotic resistance.

### The PATHOgenex RNA Atlas Provides Opportunities for Exploring Gene Expression in Human Pathogens

All transcription data have been collected in an interactive database designed to serve researchers, the PATHOgenex RNA atlas (www.pathogenex.org). The data can be browsed by selecting the strain of interest and searched with one or multiple locus tags, protein ID, gene product, and PGFam groups. PATHOgenex provides many opportunities for exploring regulation and function of pathways and gene products in human pathogens.

As example, we here show differential expression of type VI secretion systems (T6SSs) in *P. aeruginosa* PAO1. The genes encoding effectors and other components of T6SS are organized in operons, and many bacteria harbor multiple T6SS operons. Different operons are thought to be regulated differently and have different functions, but current knowledge about their regulation is limited. Utilizing PATHOgenex, we can now retrieve information of their regulation, supplying a basis for further studies. PATHOgenex retrievals show that the three *P. aeruginosa* PAO1 T6SS operons are regulated under different stress conditions. T6SS-1 is induced upon exposure to acidic and oxidative stress and to some extent upon nitrosative and osmotic stress but repressed in hypoxia and stationary phase conditions. T6SS-2 is not induced by oxidative stress and repressed both by temperature raise and during stationary phase, T6SS-3 on the other hand is induced during stationary growth, implying a different function and regulation for these effectors **(Figure 7A)**.

**Figure 7.**
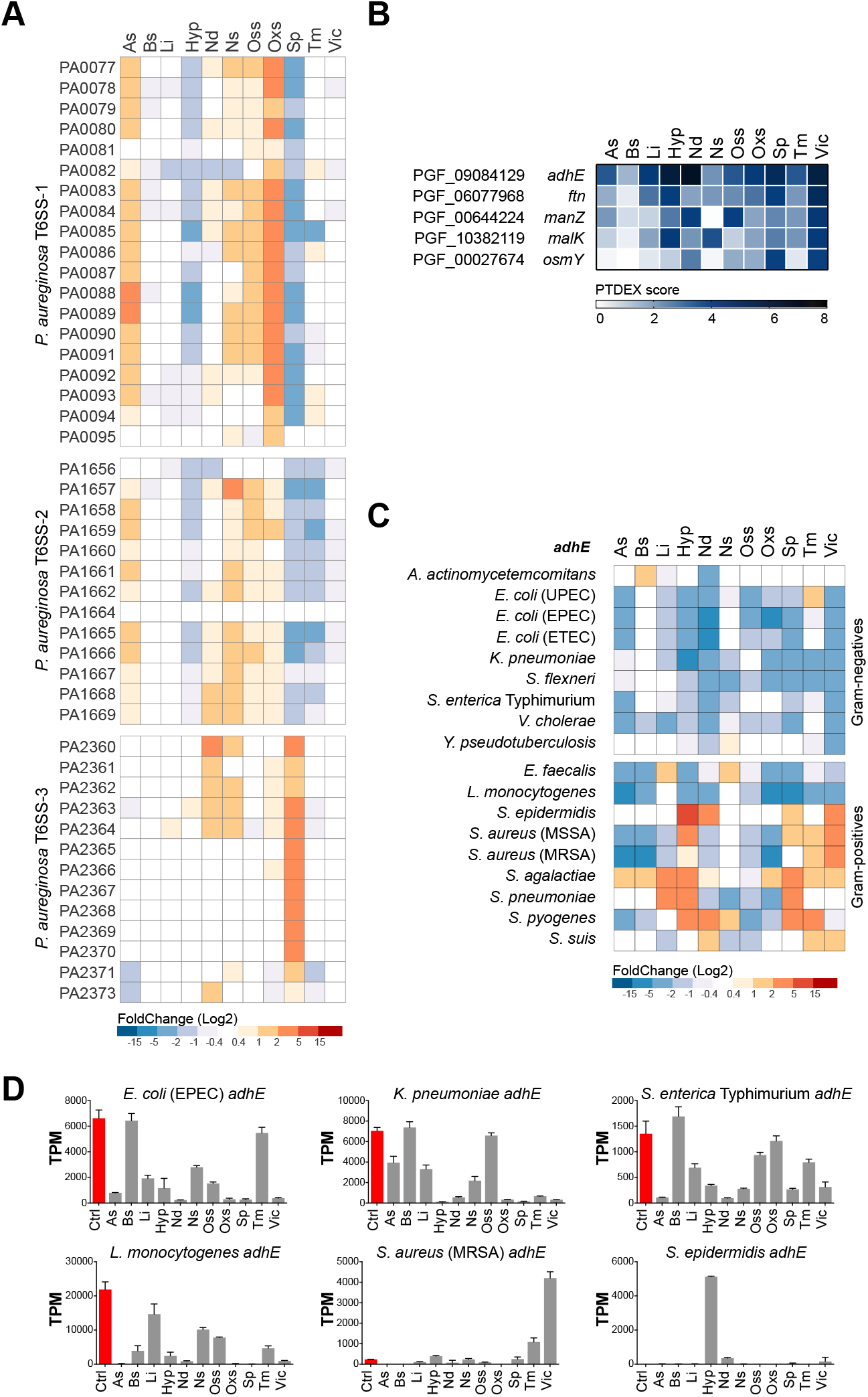
PATHOgenex RNA atlas provides opportunities for retrival of gene expression data including differential expression values, PTDEX scores and gene expression values. **(A)** Differential expression of the three T6SS operons in *P. aeruginosa* PAO1 under 11 stress conditions. **(B)** Top 5 PGFam groups with highest PTDEX scores in virulence inducing condition. **(C)** Differential expression of *adhE* in all strains carrying the gene. **(D)** Expression levels (TPM) of *adhE* across all conditions in three strains representative of Gram-negative and -positive strains. The differential expression in (A) and (C) are shown as log2 fold change with FDR-corrected *p-value* < 0.05.

PATHOgenex also provides opportunities for identification of gene products regulated in responses to certain conditions by utilizing condition-specific PTDEX scores. As an example, when retrieving the 5 PGFam groups with highest PTDEX scores in virulence inducing condition, the bifunctional acetaldehyde dehydrogenase/alcohol dehydrogenase *adhE* appeared in the top **(Figure 7B)**. This gene product is known to be involved in bacterial metabolism, but has also been attributed other functions such as stress associated translational control and virulence (Beckham et al., 2014; Kurylo et al., 2018). PTDEX scores of *adhE* and others can be analyzed further by retrieving information on which bacteria have this gene and how this gene is regulated in the different species **(Figure 7C)**. The degree of fold-change is important to know for data assessment as small fold changes in highly expressed genes are expected to have a greater impact than small changes in genes with low expression. Therefore, PATHOgenex also offers possibilities for retrieving information of expression levels for individual species (**Figure 7D)**.

Taken together, PATHOgenex provides comprehensive information of gene expression at the gene level in a particular species, but also to a broader level including gene groups and expression in multiple species. All of this information provides new opportunities for researchers to generate novel hypotheses and design experiments accordingly.

## Acknowledgments

We are grateful to all the expert labs for opening their lab to us and help with bacterial culturing and stress exposure experiments. We thank Drs. Peter Lind, Teresa Frisan, and Saskia Erttmann for critical reading of the manuscript. The work was supported by Knut and Alice Wallenberg Foundation (No. 2016.0063), Swedish Research Council (No. 2018-02855), and Insamlingsstiftelsen, Medical Faculty at Umeå University to M Fallman; Novo Nordisk Foundation (K. Avican was partly supported by grant, No. NNF17OC0026486, awarded to Dr. Emmanuelle Charpentier at MIMS, The Laboratory for Molecular Infection Medicine Sweden); ERC (Starting grant, No. 716063) to J. Tang; Academy of Finland Research Fellow Grant (No. 317680) to J. Aldahdooh.

## Author Contributions

K.A. and M.F. conceived and supervised the project, and wrote the manuscript with input from J.A, M.T., J.T., K.B., and M.R.. K.A. performed the experiments and analysed the data. M.T., K.B., K.A, and M.F. designed and M.T. calculated PTDEX scores. J.A., J.T., K.A., and M.F. designed and J.A. constructed the PATHOgenex RNA atlas and webpage.

## METHOD DETAILS

### Bacterial growth and stress exposures

All bacterial strains shown in **Figure 1A** were grown in their optimal growth medium and temperatures as indicated in **Table S1** in laboratories specialized in each species. Three bacterial cultures were grown overnight and then subcultured to a new culture vial with 1:50-1:100 dilution. The cultures were grown until exponential phase with an OD_600_ of 0.1-0.6 and exposed to the 11 stresses separately. The stress conditions, reagents used to generate them, and the exposure times are given in **Table S1**. Unexposed cultures at exponential growth were used as controls for differential expression analysis. For nutritional downshift, bacterial cultures were spun down at 9000 rpm for 2 minutes, the supernatant removed, the pellets resuspended in 1X M9 supplemented with 0.1 M MgCl_2_ and 0.1 M CaCl_2_, and incubated for 30 minutes at ambient temperature for each strain. For hypoxia, 2 ml screw cap tubes were filled with bacterial culture at exponential growth and the caps were screwed on without leaving space for air. The stress exposures were stopped by adding 5% (final concentration) phenol:ethanol solution **(Figure S1)**.

### Total RNA isolation

One milliliter of triplicate bacterial cultures exposed to the stresses and un-exposed control culture were immediately pelleted by centrifugation at 9000 rpm at room temperature for 2 minutes after adding phenol:ethanol. The supernatants were removed and pellets resuspended in 0.5 ml Trizol solution. For Gram-negative bacteria, cells were homogenized in Trizol solution by pipetting up and down 15 times. For Gram-positive bacteria, culture suspensions in Trizol were transferred to previously cooled bead beater tubes containing 0.1-mm glass beads and treated with Mini-Beadbeater (Biospec Products Inc, USA) twice at a fixed speed for 45 seconds, and then cooled on ice for 1 minute between the treatments. Culture homogenates were incubated at room temperature for 5 minutes and then supplemented with 0.2 ml chloroform, thoroughly mixed by shaking 10 times, and incubated for 3 minutes. After centrifugation at 12 000 g at 4°C for 15 minutes, the aqueous upper phase was carefully transferred to new RNase free tubes. An equal volume of 99% ethanol was added to the aqueous phase and isolation continued using the Direct-Zol RNA Miniprep Plus (Zymo Research, USA) RNA purification kit protocol. Total RNAs were eluted in RNase free water in RNase free tubes. The total RNA concentrations were measured using the Qubit BR RNA Assay Kit (ThermoFisher Scientific, USA) and RNA integrity confirmed on a 0.8% agarose gel in TBE buffer.

### RNA-seq library preparation with RNAtag-Seq

All rRNA-depleted RNA-seq library preparations were performed according to Shishkin et al. (2015), with minor modifications. RNAtag-Seq allows multiple library preparations in one tube, with initial tagging of total RNA samples with modified (5’P and 3’ SpcC3) DNA barcoded adaptors, each harboring a unique 8 nt sequence used to demultiplex individual libraries after sequencing. We combined the library preparation of three biological replicates from 11 stress-exposed samples and the un-exposed control sample, resulting in 36 library preparations in one tube for each bacterial strain. To tag the 36 replicates, 36 unique barcoded adaptors were used, with every three barcodes used for samples from the same stress conditions in all bacterial species **(Table S2)**. A total of 100 ng total RNA was used for each biological replicate. The total RNA was fragmented in 2X FastAP Thermosensitive Alkaline Phosphatase buffer for 3 minutes at 94°C, DNase treated, and dephosphorylated with a combination of TURBO DNase and Thermosensitive Alkaline Phosphatase in 1X FastAP buffer for 30 minutes at 37°C. Fragmented, DNase-treated, and dephosphorylated total RNAs were cleaned with a 2X reaction volume of Agencourt RNAClean XP beads. Cleaned total RNAs were incubated with 100 pmol of the unique DNA barcode adaptors at 70°C for 3 minutes, and then ligated with T4 RNA ligase 1 for 90 minutes at 22°C. The ligation was stopped and the enzyme denatured by the addition of RLT buffer. The 36 denatured ligation mixes were then pooled and cleaned in a Zymo Clean & Concentrator™-5 column according to the manufacturer’s 200 nt cut-off protocol. The RNAs were eluted in 32 μl of RNase free water. Ribosomal RNA was depleted using the Ribo-Zero™ Magnetic Gold Kit (Bacteria) according to the manufacturer’s instructions. The first strand cDNA of each pool was generated using an AffinityScript Multiple Temperature cDNA synthesis kit with 50 pmol of AR2 primer at 55°C for 55 minutes. The RNA was degraded by adding a 10% reaction volume of 1 N NaOH at 70°C for 12 minutes and the reaction neutralized with an 18% reaction volume of 0.5 M acetic acid. After cleaning the reverse transcription primers with a 2X reaction volume of RNAClean XP beads, the 3Tr3 adapter was ligated with T4 RNA ligase 1 at 22°C with overnight incubation. The second ligation was cleaned first with a 2X, and secondly 1.5X, reaction volume of RNAClean XP beads. The cDNA was then used as the template for PCR reaction with FailSafe™ PCR enzyme mix using 12.5 pmol 2P_univP5 as forward primers and 12.5 pmol ScriptSeq™ Index (barcode) PCR primers as reverse primers. The PCR cycles were as follows: 95°C for 3 minutes, followed by 12 cycles at 95°C for 30 seconds, 55°C for 30 seconds, and 68°C for 3 minutes, and then finishing at 68°C for 7 minutes. The PCR product was cleaned first with a 1.5X, and secondly 0.8X, reaction volume of AMPure beads and eluted in RNAse free water. The library concentrations were measured using the Qubit™ dsDNA HS Assay Kit and the library insert size determined by the Agilent DNA 1000 Kit in a 2100 Electrophoresis Bioanalyser Instrument (Agilent, USA).

## QUANTIFICATION AND STATISTICAL ANALYSIS

### RNA-seq data analysis

The RNA-seq libraries generated by RNAtag-Seq were sequenced by either single end or paired end Illumina sequencing at SciLifeLab, Stockholm. Each library harboring a pool of 36 RNA samples was demultiplexed according to the unique 8 nt on the ligated barcode adaptors to separate reads from each replicate. The sequencing reads generated in this study were deposited in GEO with accession number GSEXXXXXX. Demultiplexed reads were then mapped to the genome of the species to which the RNAs belong in an annotation-independent manner. The accession numbers of the RefSeq reference genomes used for each strain can be found in “Microbes and Viruses” in Key Resouces. The number of reads mapped to CDSs, rRNA, and tRNA was calculated according to the annotation files. The number of reads mapped to the non-coding regions were calculated by subtracting the reads mapped to the annotated regions from the total number of mapped reads. The sequencing reads from primary transcripts of *K. pneumoniae* (Kim et al., 2012) with accession number SRR408498 were downloaded from Sequence Read Archive (SRA) and mapped to the same reference genome used in this study. *In vivo* and *in vitro* RNA-seq reads of *S. aureus* strain 6850 (Szafranska et al., 2014) with accession number ERP005459 were downloaded from SRA and mapped to RefSeq reference genome CP006706. All of the demultiplexing and read mapping steps were performed in CLC Genomic Workbench (Qiagen, USA).

For differential gene expression analysis, the trimmed mean of M values (TMM) (Robinson and Oshlack, 2010) normalization method was used to normalize the sequencing depth of the individual libraries. For each bacterial species, the comparisons were performed between multiple groups and the control sample so that the full dataset could be used for fitting the generalized linear model. Using the complete datasets with multiple group comparisons allowed us to determine whether a gene has unstable expression for which the variation is random or is differentially expressed in response to the stress. Read values for genes with a maximum group mean expression (the maximum average TPM value in the statistical comparison group) <20 were removed and a threshold of 1.5-fold change with FDR-adjusted *p*-value <0.05 was employed to determine differential expression.

### Generation of a phylogenetic tree of 32 bacterial strains

A phylogenetic tree of the 32 strains based on NCBI taxonomy was generated with PhyloT in Newick format. The visualization of the tree was achieved using iTol.

### Clustering 32 bacterial strains based on expression profiles

Hierarchical clustering, with Euclidian distance, of 32 strains based on the percent of genes expressed (TPM≥10) in all 12 conditions, percent of genes expressed in at least one condition, and percent of genes not expressed in any of the conditions was performed with ClustVis (Metsalu and Vilo, 2015).

### Clustering 105 088 genes based on functional orthologs and isofunctional homologs

The KEGG orthology group was assigned for clustering genes based on functional orthologs. The amino acid sequences of all CDSs for each strain were uploaded to GhostKOALA (Kanehisa et al., 2016) to assign the best KO group in the genus_prokaryotes KEGG GENES database. Clustering based on isofunctional homologs was performed with PATRIC’s Proteome Comparison Service. The amino acid sequences of all CDSs for each strain were compared using its own annotated genome, if available, or the closest strain to assign the best PGFam group for each CDS.

### PTDEX score calculation

The PTDEX score was calculated for the orthology/homology groups using Python scripts and Jupyter notebooks, relying on the open-source libraries NumPy and Pandas. The PTDEX score formula relies on the definition of differentially expressed genes given in “RNA-Seq Analysis” above and the equation shown in Figure 2B, where *n_genes_(on)* is the number of differentially expressed genes in the orthology/homology gene group, *ngenes(total)* is the total number of genes in the group, *n_strains_(on)* is the number of strains for which at least one of the orthology group genes is differentially regulated, *n_strains_(total)* is the total number of strains for a given stress condition, and *n_strains_(database)* is the total number of strains in the database (*n_strains_(database)* =32 for general PTDEX scores, 21 for Gram negative-specific, 9 for Gram-positive-specific PTDEX scores). The additional logarithmic component is a weight factor to give more relative importance to orthology/homology groups with more genes.

### Generation of co-expression modules on PGFam PTDEX scores and associated KEGG pathway enrichments

The co-expression Modules Identification tool (CEMiTool) (Russo et al., 2018) was implemented on PGFam groups of Gram-negative and -positive strains to find PGFam groups with similar PTDEX score patterns over the 11 stress conditions. PGFam groups associated with each module can be found in Table S5 and Table S6. Each PGFam group associated with the particular module, if possible, was converted in KO groups and KO groups were used in the KEGG orthology database to map pathways.

### PATHOgenex website construction

PATHOgenex was built using PHP Laravel Framework 6.18.2 and PHP 7.4.4 for server-side data processing, Javascript ECMAScript 2015 for the front end, and D3.js 5.16.0 and Plotly 1.40.0 libraries for the generation of the interactive visualizations. Linux distribution CentOS-8 with the 64-bit kernel 4.18.0 running on four processor cores and 64 Gb of RAM is used to host the web service on the in-house computational cluster.

---

**Figure S1. Stress exposure and experimental setup. Related to Figure 1**. Three bacterial cultures were inoculated from bacteria grown in the stationary phase for the stress exposure experiments. Cultures, except control samples, were exposed to the stress conditions at exponential growth (OD_600_ 0.1-0.5). Transcription was stopped by adding 0.05% (final concentration) phenol:ethanol solution. Cells were pelleted and homogenized in Trizol solution and stored at -80 °C until RNA extraction step. The libraries were generated with RNAtag-Seq method from total RNA. The specific culturing and stress conditions for each species are indicated in **Table S1**. *L. pneumophila* and *M. tuberculosis* were not exposed to the virulence inducing condition, no such condition has been described.

**Figure S2. The global expression pattern of biological replicates from the same sample cluster. Related to Figure 2.** Correlation coefficients (*R*) of the global expression levels were plotted on a matrix and a dendogram showing the clustering of replicates with complete linkage (below) for each species. *R* values were calculated by the Pearson method. The matrices were generated using the “corrplot” R package and complete linkage by the CLC Genomics Workbench.

**Figure S3. Differential expression of genes previously known to be regulated under particular stress conditions and percentage of differentially expressed genes in different groups of bacteria. Related to Figure 3. (A)** Differential expression of; *bfd* encoding bacterioferritin-associated ferredoxin, in low iron conditions, *fnr* encoding fumarate and nitrate reductase under hypoxia, *cstA* encoding carbon starvation protein under nutritional downshift, *groL* encoding heat shock protein under high temperature conditions, *dps* encoding DNA protection during starvation in the stationary phase, *ahpC* encoding alkyl hydroperoxide reductase C under oxidative stress conditions, *proW* encoding glycine betaine/proline transport system permease protein under osmotic stress conditions, *hmp* encoding flavohemoprotein, which is involved in NO detoxification, under nitrosative stress conditions. **(B)** Aerobic (gray) and microaerophilic bacteria (orange). **(C)** *Bacilli* (gray), *Betaproteobacteria* (orange), and *Gammaproteobacteria* (red). **(D)** C1 bacteria (gray) and C2 bacteria (orange) identified in Figure 2D. Error bars indicate standard deviation of the mean. The significance between the groups was calculated with Multiple t-test using Holm-Sidak method. ** indicates adjusted *p*-value < 0.01.

**Figure S4. Diverse transcription factors from Gram-negative and Gram-positive bacteria shows disctinct PTDEX score patterns in bile stresses. Related to Figure 5.** PTDEX scores of sigma factors, response regulators, and one-component systems in both Gram-negative and -positive strains. Transcription factors in the different bacteria were identified using the P2TF database (Ortet et al., 2012).

**Figure S5. PTDEX score ≥0.25 is highly accurate for indicating the probability of gene groups being regulated under a particular stress condition. Related to Table 1.** Randomly picked three PGFam groups from Gram-negative strains with PTDEX scores between 0.25-0.3 and different numbers of genes included for each stress condition to show how the PTDEX score indicates the differential expression of the gene groups. Each box represents an individual gene from the gene group. Red indicates downregulation, blue indicates upregulation, white indicates not regulated, and crossed white indicates a gene for which expression could not be measured due to lack of RNA.

**Figure S6. PTDEX scores identify universal stress responder genes involved in responses to multiple stresses in both Gram-negative and -positive strains. Related to Table 1.** USRs were clustered in different biological processes using the PATRIC Subsystems tool and are shown with their general PTDEX scores over the 11 stress conditions.

**Figure S7. Sequence homology of KPN_01149 paralogs in *K. pneumoniae* and expression of the intergenic region in *S. aureus* from *in vivo* samples. Related to Figure 6. (A)** Multiple sequence alignments of KPN_01149 and its paralogs in *K. pneumoniae*. Sequences marked with blue indicate the 5’-UTR and 3’-UTR. Red-colored nucleotides indicate identical nucleotides among the three sequences. **(B)** Reads mapped to *S. aureus* 6850 intergenic region described in Figure 6G during acute infection and chronic infection in a murine infection model, and *in vitro* exponential growth (Szafranka et al., 2014).

