## Supplementary figures and images for "RNA Atlas of Human Bacterial Pathogens Uncovers Stress Dynamics Linked to Infection"

### Supplementary Figure 1

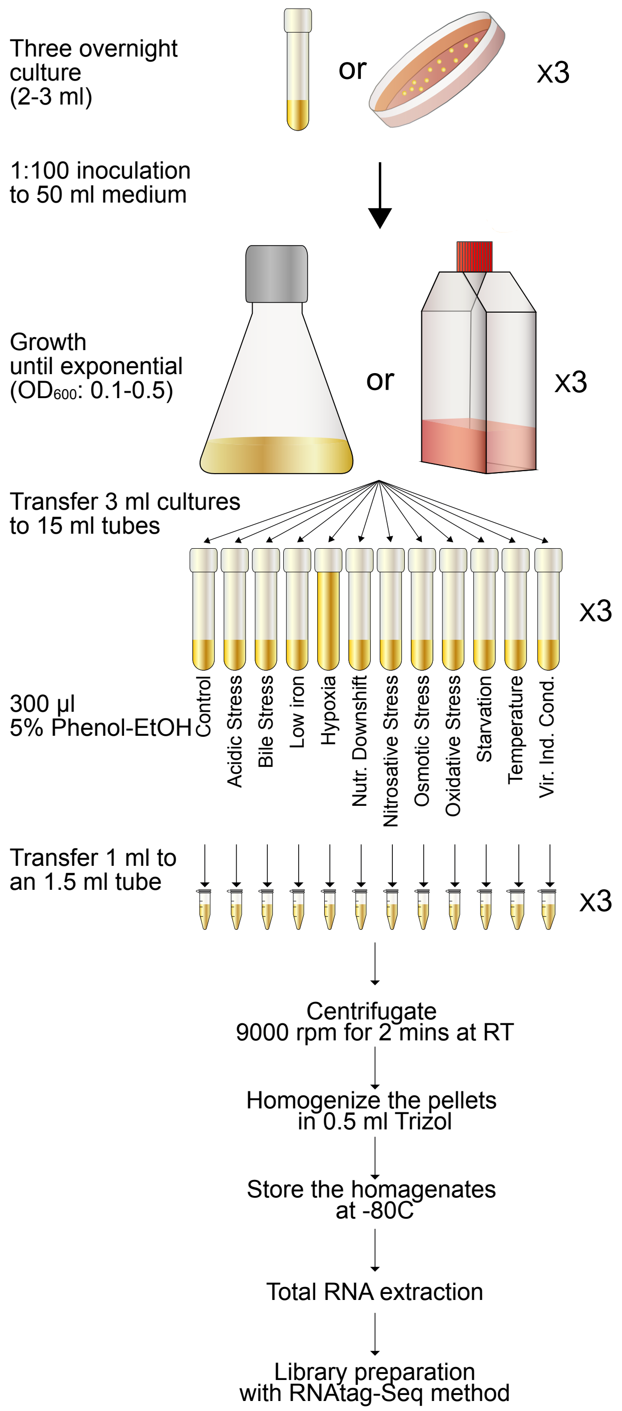

### Supplementary Figure 2

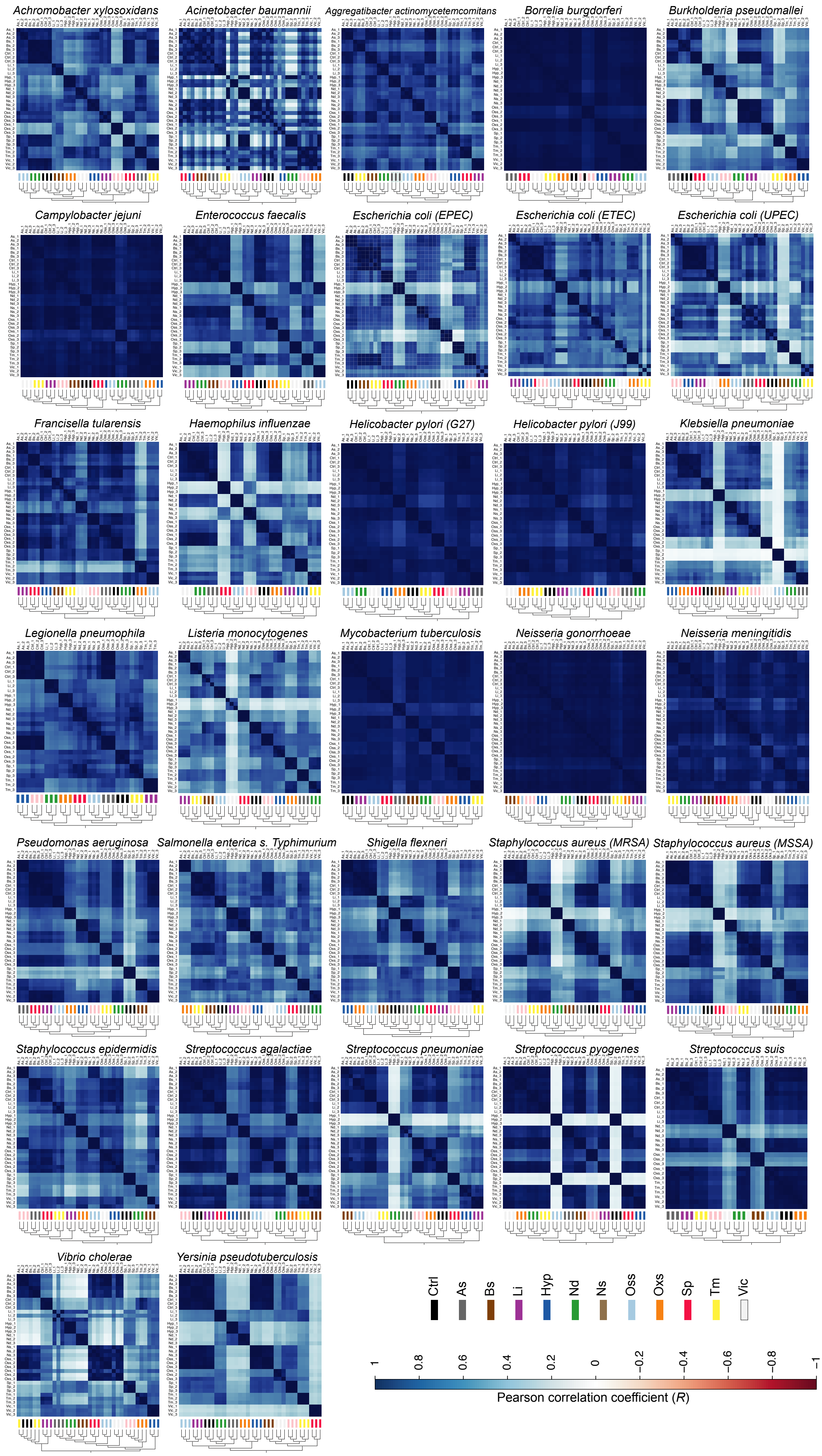

### Supplementary Figure 3

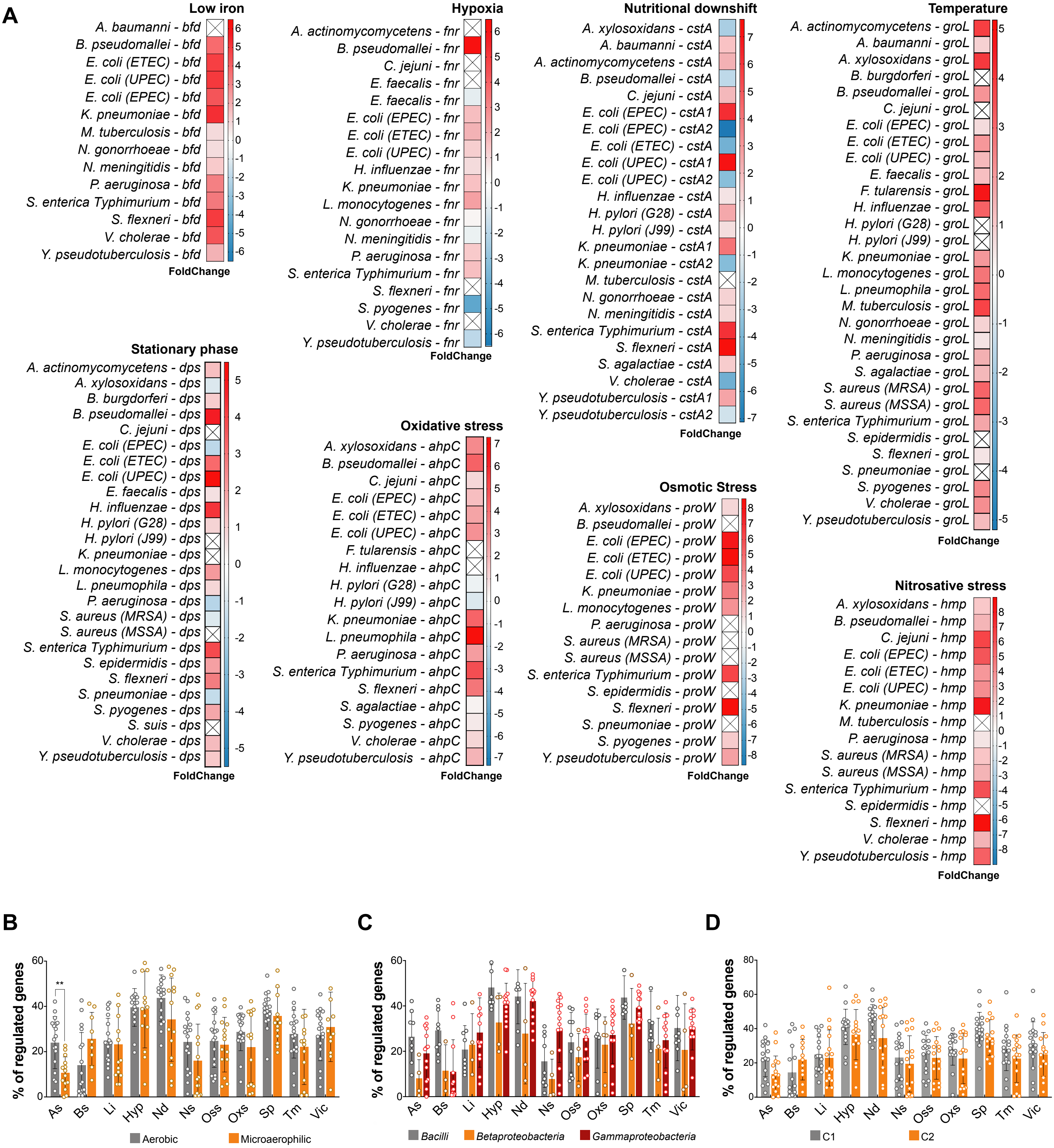

### Supplementary Figure 4

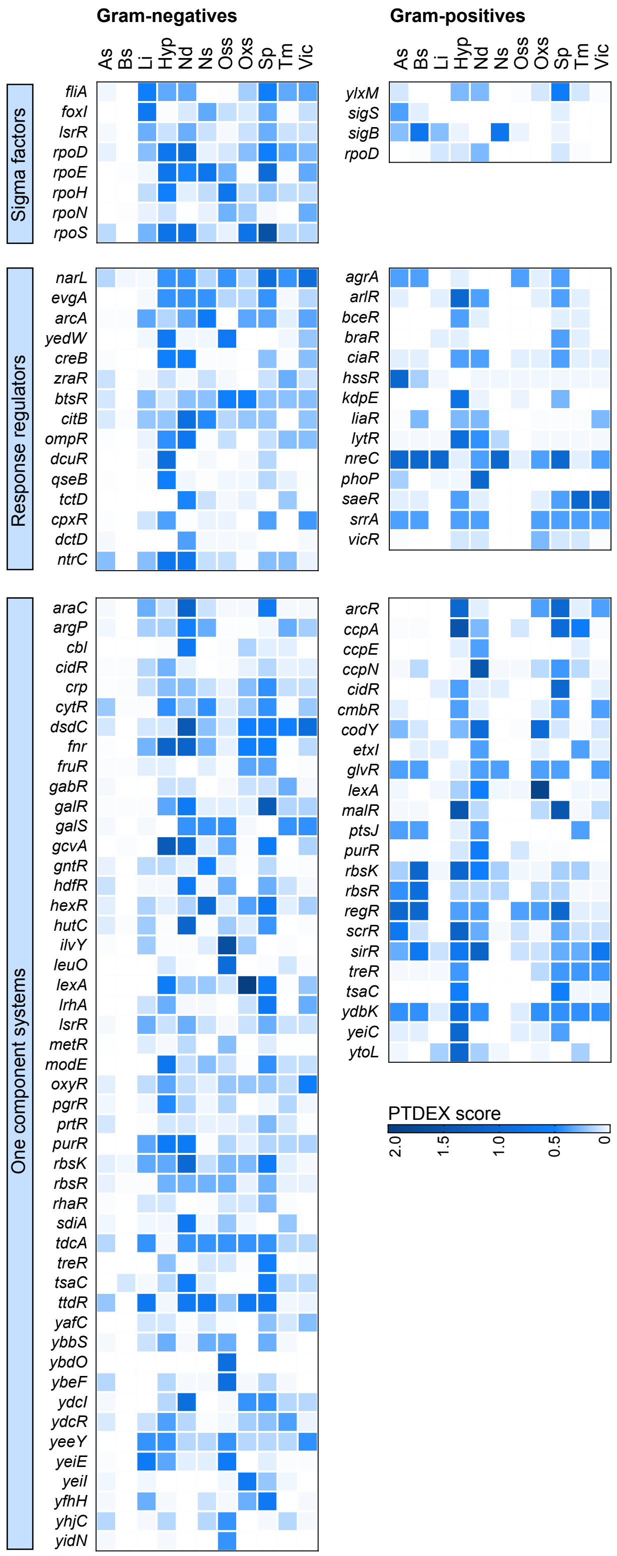

### Supplementary Figure 5

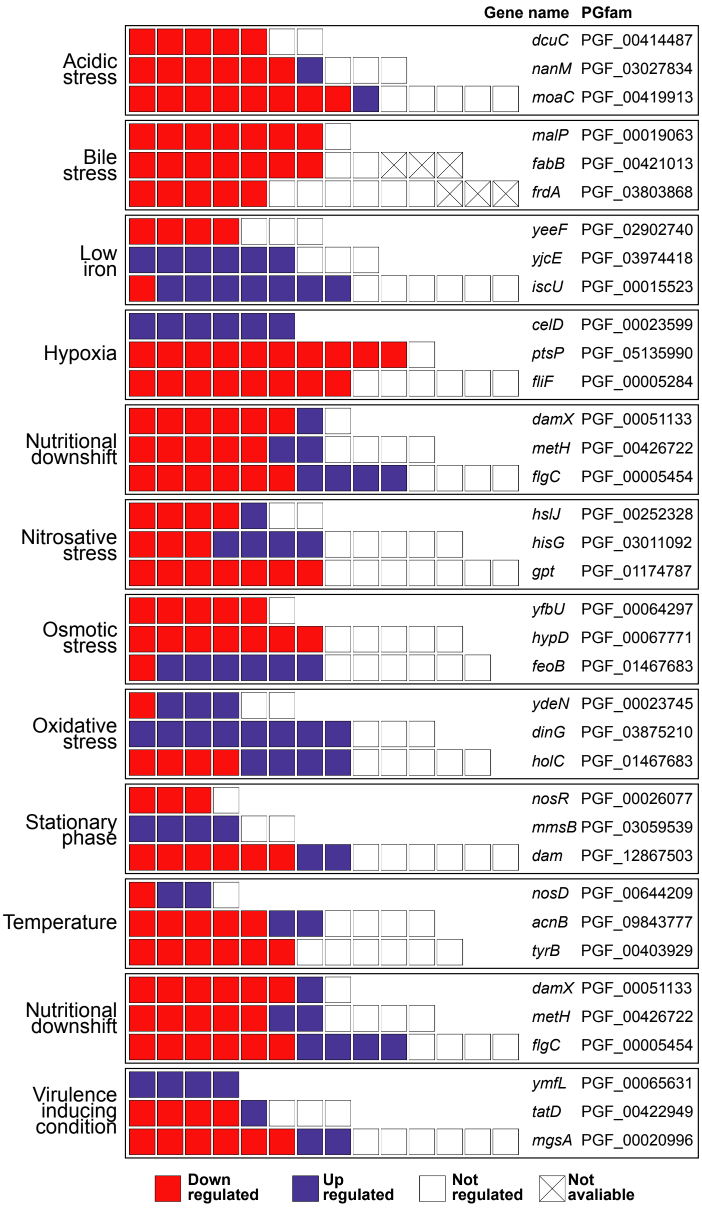

### Supplementary Figure 6

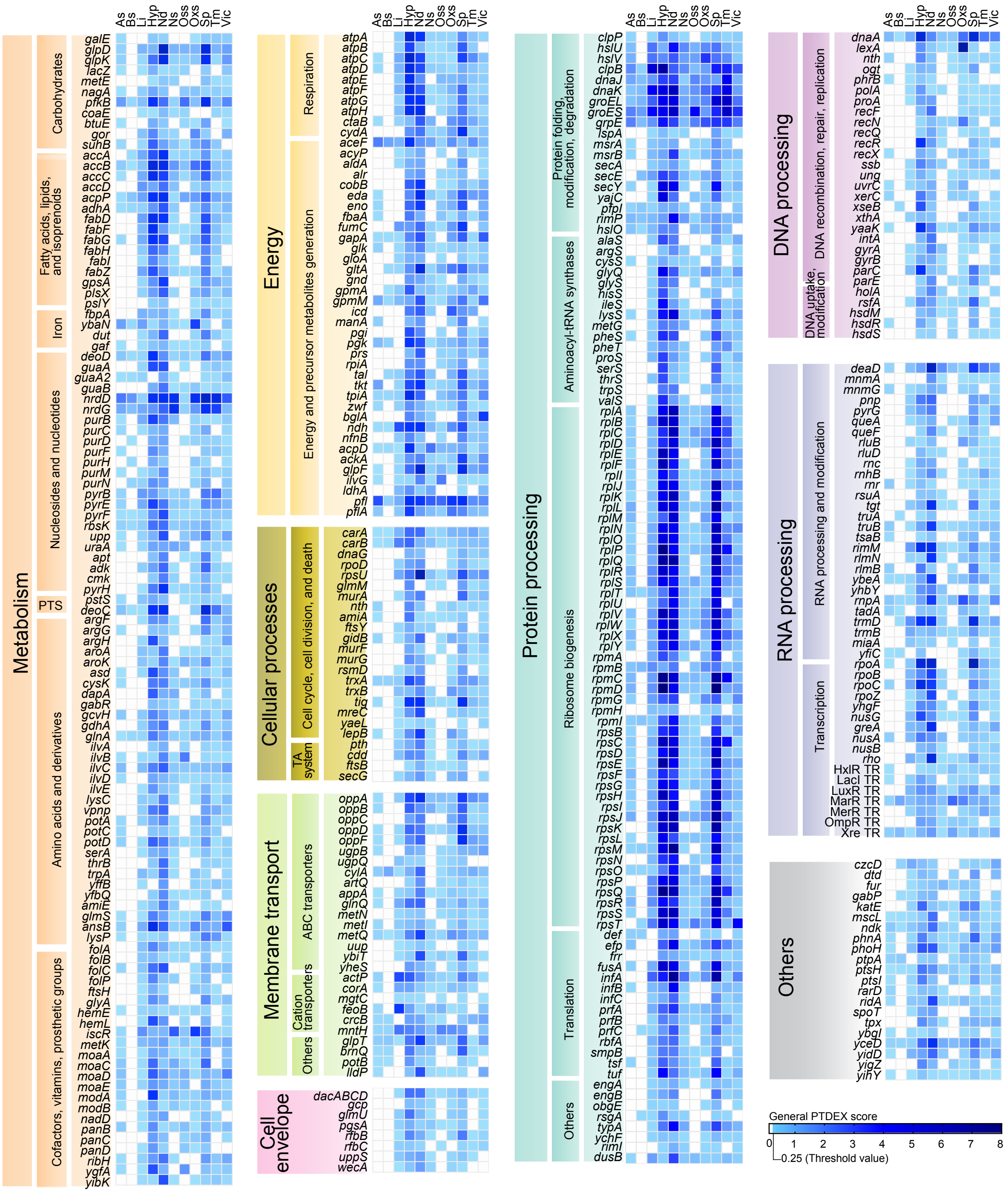

### Supplementary Figure 7

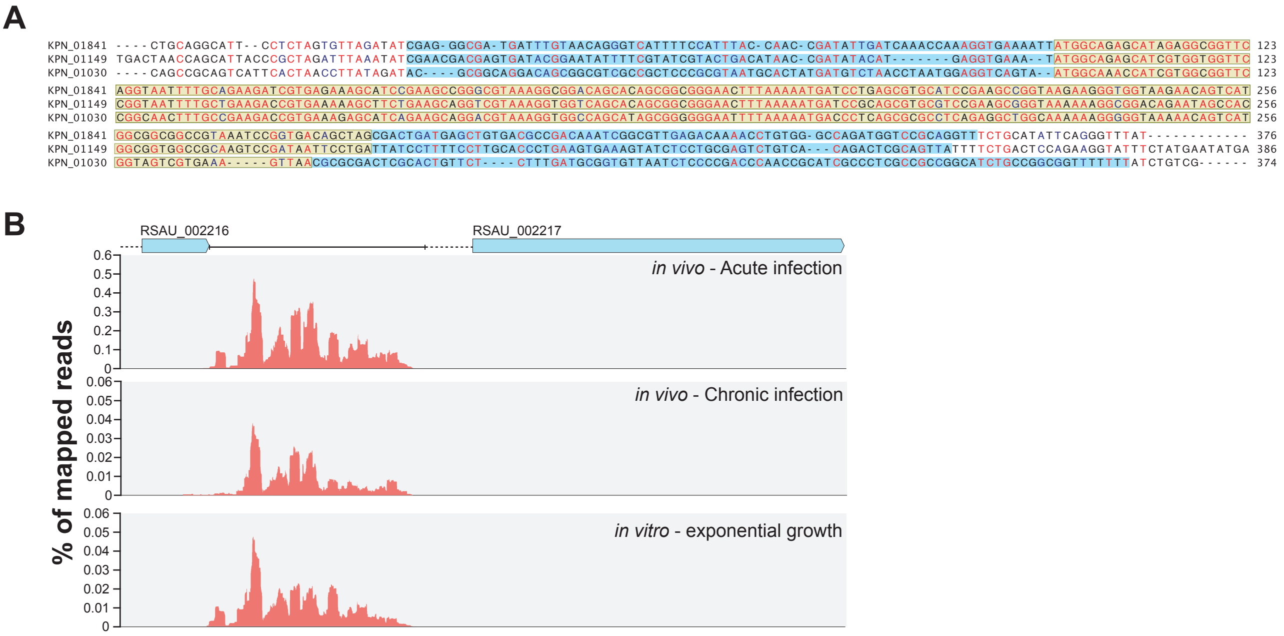
